# FGFR1-phosphate sensing and crystal-induced gasdermin D signaling in neutrophils drive vascular calcification in CKD

**DOI:** 10.64898/2026.05.23.725305

**Authors:** Atsuko Miyoshi-Harashima, Daigo Nakazawa, Tetsuo Shimizu, Kanako Watanabe-Kusunoki, Fumihiko Hattanda, Saori Nishio, Satoka Shiratori-Aso, Yusho Ueda, Mizuki Kimura, Takuro Kawamura, Shun Takenaka, Masatoshi Kanda, Sakiko Masuda, Yuka Nishibata, Utano Tomaru, Yasushige Shingu, Satoru Wakasa, Akihiro Ishizu, Tatsuya Atsumi

**Affiliations:** Department of Rheumatology, Endocrinology, and Nephrology, Faculty of Medicine and Graduate School of Medicine, Hokkaido University, Sapporo, Japan; Division of Rheumatology and Clinical Immunology, Sapporo Medical University, Sapporo, Japan; Department of Medical Laboratory Science, Faculty of Health Sciences, Hokkaido University, Sapporo, Japan; Department of Surgical Pathology, Hokkaido University Hospital, Sapporo, Japan; Department of Cardiovascular Surgery, Faculty of Medicine and Graduate School of Medicine, Hokkaido University, Sapporo, Japan

## Abstract

Chronic kidney disease (CKD) confers disproportionate cardiovascular risk. In non-dialysis CKD, calcification accumulates primarily within the intimal layer. Clinical studies indicate that intimal calcification correlates with hyperphosphatemia, yet the cellular and molecular pathways remain unclear. Given evidence that osteocytes sense phosphate via fibroblast growth factor receptor 1 (FGFR1), we hypothesized that FGFR1-expressing vascular immune cells, especially neutrophils, act as mediators linking high phosphate to plaque mineralization. In vitro, phosphate triggered FGFR1-dependent signaling in human and murine neutrophils, inducing neutrophil extracellular traps (NETs). Activated neutrophils promoted the depletion of Fetuin-A, a major inhibitor of calcium–phosphate complexation, creating a milieu permissive to mineral nucleation. Newly formed calcium–phosphate particles amplified NETs through gasdermin D (GSDMD), establishing a feed-forward loop that enhanced mineralization and endothelial injury in co-culture assays. Human arteriosclerotic plaques from CKD patients showed NETs markers co-localizing with calcified deposits. In vivo, pharmacological FGFR inhibition attenuated arterial intimal calcification and suppressed NET formation in CKD mice. These findings identify phosphate sensing via neutrophil FGFR1 and subsequent crystal-induced GSDMD signaling as drivers of intimal vascular calcification in CKD. Targeting phosphate-sensing pathways, NET formation, and neutrophil-driven mineralization may mitigate vascular calcification and reduce cardiovascular risk in CKD.

## Introduction

Chronic kidney disease (CKD) is a progressive condition characterized by a high risk of cardiovascular disease (CVD). Even in the early stages of CKD, the prevalence of fatal CVD events is elevated, with arteriosclerosis and vascular calcification emerging as critical pathological features (1–4). Notably, in non-dialysis CKD patients, the calcification primarily occurs within the intimal layer of the vasculature, in contrast to the medial calcification often seen in end-stage renal disease patients undergoing dialysis (5, 6). Although higher serum phosphate concentrations have been associated with a greater prevalence of intimal vascular calcification in CKD (7), phosphate-lowering agents appear to have limited efficacy in controlling intimal calcification (8, 9), and the underlying mechanisms remain poorly understood. Recent research has uncovered elements of the phosphate-sensing system, showing that osteocytes sense elevated serum phosphate through FGFR1, thereby influencing phosphate homeostasis (10). Among vascular and immune cells, neutrophils have been identified as a key population expressing FGFR1 (11). Given their established role in atherosclerotic plaque formation—where neutrophils respond to oxidized low density lipoprotein (LDL), producing reactive oxygen species (ROS) and releasing NETs (12, 13), we hypothesized that neutrophils might similarly react to elevated phosphate levels in CKD, potentially promoting intimal calcification. Recent studies have shown that pyroptosis in neutrophils contributes to NETosis by activating GSDMD, which forms membrane pores that enable calcium influx and activation of peptidylarginine deiminase 4 (PAD4), which catalyzes histone citrullination to enable chromatin decondensation and NET release (14). Moreover, crystalline materials, including calcium phosphate crystals, can also induce NETs through GSDMD-dependent pathways, potentially promoting intimal calcification (15, 16). We explored the role of neutrophils in phosphate-driven calcification and vascular injury in CKD.

## Results

Neutrophils from healthy donors were cultured in commercially available DMEM (containing baseline Ca 1.8 mM and P 0.9 mM) supplemented with 2.5% FBS and additional phosphate to achieve varying concentrations. NET formation was assessed by SYTOX Green staining (Figure 1A). Neutrophils formed SYTOX-positive NETs, with significantly larger SYTOX-positive areas under high phosphate conditions (Figure 1, B and C). Time-lapse imaging under high phosphate conditions showed a progressive increase in NET formation (Figure 1, D and E).

**Figure 1.**
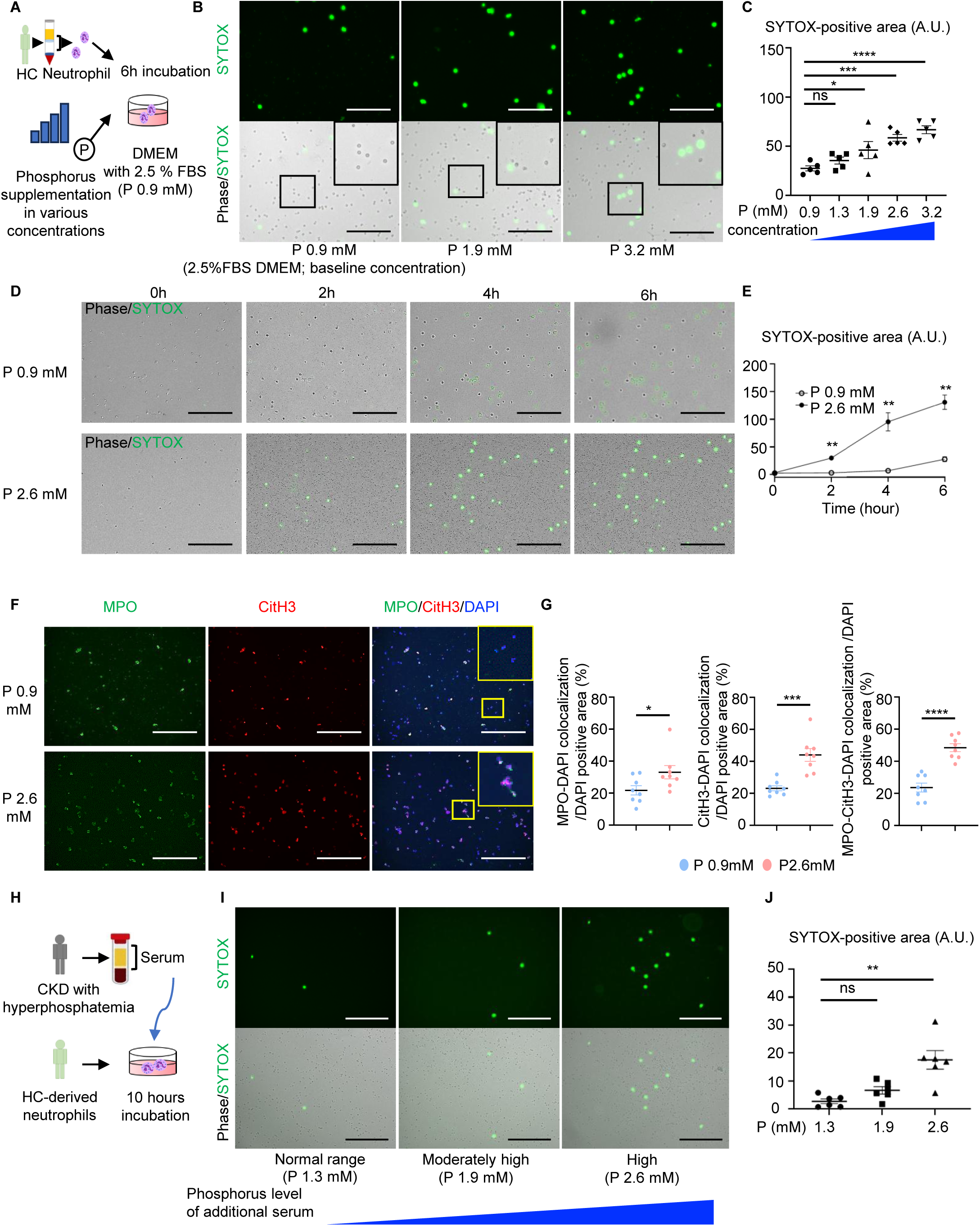
Neutrophil activation in response to elevated phosphate concentration. (A) Peripheral blood neutrophils from healthy controls (HC) (1×10^5^ cells/ml) were incubated for 6 hours in DMEM supplemented with 2.5% FBS and varying phosphate (P) concentrations. Note that the baseline phosphate concentration (0.9 mM) is intrinsic to commercially available DMEM and was not additionally supplemented. Cells were then stained with SYTOX green. (B) The representative immunofluorescence images of SYTOX green (green) staining. (C) The quantification of SYTOX-positive area. n=5. (D) The representative time-lapse images of neutrophils stimulated by high or low P concentrations. (E) The quantification of SYTOX-positive area every 2 hours. n=3. (F) The representative immunofluorescence images of MPO (green), CitH3 (red) and DAPI (blue). (G) The quantitative analysis of neutrophils for the MPO-DAPI double-positive area per DAPI positive area, the CitH3-DAPI double-positive area per DAPI positive area, and the MPO-CitH3-DAPI triple-positive area per DAPI positive area, respectively. n=8. (H) The serum from CKD patients with normal phosphate levels (P 1.3mM), moderately high phosphate levels (P 1.9mM), and high phosphate levels (P 2.6mM) was added to healthy neutrophils and evaluated. (I and J) The representative images and a comparison of the quantification of SYTOX-positive area. Data represent mean ± SEM. Statistical analysis was performed using one-way ANOVA with post hoc Dunnett’s multiple-comparison test (C and J) or multiple t test (E) or Student’s unpaired t test (G). *P <0.05, **P < 0.01, ***P < 0.001, ****P < 0.0001. ns: not significant. The insets are higher magnification images of the black and yellow boxes. Scale bars: 200 μm (B, D); 100 μm (F and I).

Immunofluorescence for CitH3 (a key NET marker) and myeloperoxidase (MPO, a neutrophil marker) confirmed the formation of MPO-CitH3-DAPI-positive NETs under high phosphate (Figure 1, F and G). We next compared NET formation between neutrophils from healthy controls (HC) and pre-dialysis CKD patients (Supplemental Figure 1A), whose characteristics are detailed in Supplemental Table 1. CKD neutrophils formed NETs even at lower phosphate levels, with significantly larger SYTOX-positive areas than HC (Supplemental Figure 1, B and C), indicating heightened sensitivity. To investigate the actual impact of serum phosphate levels in CKD patients on neutrophils, serum from CKD patients with varying phosphate levels was added to healthy neutrophils and incubated for 10 hours (Figure 1H). Serum with high phosphate levels significantly induced NET formation (Figure 1, I and J). In contrast, lymphocytes and monocytes did not respond to high phosphate (Supplemental Figure 2, A and B), suggesting that the response to high phosphate is specific to neutrophils.

### High phosphate concentrations induce NET formation through the activation of FGFR1 signaling pathway

We investigated the role of FGFR1 in phosphate signaling in neutrophils. Neutrophils were exposed to high phosphate with or without 20 µM FGFR1 inhibitor (SU5402), which reduced phosphate-induced NET formation (Figure 2, A and B). To exclude the possibility of cytotoxicity or off-target effects, we first confirmed that SU5402 did not affect neutrophil viability at concentrations ranging from 2.5 to 80 µM (Supplemental Figure 3). Furthermore, to genetically validate the role of FGFR1, we employed HL-60 cells differentiated with all-trans retinoic acid as a neutrophil model (17) and transfected them with FGFR1-targeting siRNA (Figure 2C). Cell viability following transfection remained intact (Supplementary Figure 4), and qPCR confirmed efficient FGFR1 knockdown (Figure 2D). Under high phosphate conditions, HL-60 cells treated with a non-targeting control siRNA formed NETs with histone citrullination, whereas FGFR1-silenced cells did not (Figure 2, E-H). Given that FGFR1 activates ERK pathway and histone citrullination is downstream of ERK-ROS activation (18), (19), we examined this pathway. High phosphate induced ERK phosphorylation in control HL-60 cells, but this was abolished by FGFR1 silencing (Figure 2, I and J). Together, these results suggest that elevated phosphate activates FGFR1–ERK signaling pathway leading to histone citrullination and NET formation.

**Figure 2.**
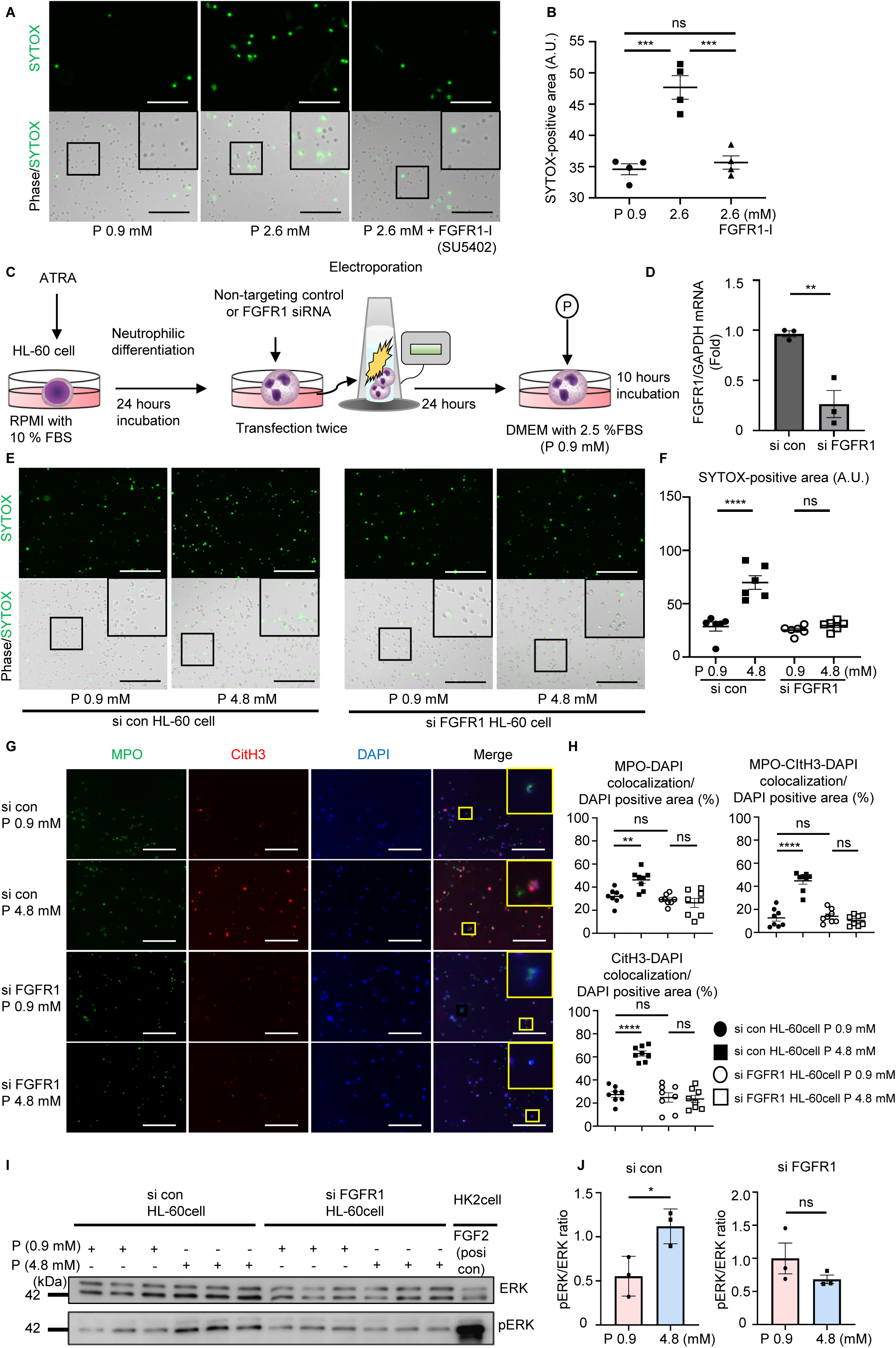
The role of FGFR1 in neutrophils as a sensor of elevated phosphate levels. (A) Neutrophils pretreated with a 20 μM FGFR1 inhibitor (FGFR1-I, SU5402) were cultured under high phosphate conditions for 6 hours, subsequently stained by SYTOX green. (B) The quantification of SYTOX-positive area. n=4. (C) HL-60 cells were differentiated into neutrophils by incubation with ATRA for 24 hours. The cells were then transfected with FGFR1-specific siRNA via electroporation, performed twice at 24-hours intervals. As a control, HL-60 cells were electroporated under the same conditions with non-targeting control siRNA. At 24 hours after the second transfection, both control and FGFR1 knockdown HL-60 cells were cultured under high phosphate conditions for 10 hours. (D) The expression of FGFR1 mRNA in differentiated HL-60 cells transfected with control or FGFR1-targeting siRNA, assessed by qPCR (n=3). (E-H) FGFR1 knockdown HL-60 cells and control HL-60 cells were incubated under 0.9mM or 4.8mM phosphate conditions for 10 hours. NET formation was assessed by (E and F) SYTOX green (green) (n=6) and (G and H) CitH3 staining (red), with MPO (green) serving as a neutrophil marker (n=8). (I) Immunoblotting was performed using antibodies against ERK and pERK on lysates derived from FGFR1 knockdown or control HL-60 cells under low or high phosphate conditions. As a positive control (posi con), human kidney-2 (HK2) cells exposed by FGF2, one of the FGFR ligands, were used. (J) The ratio of pERK/ERK expression. n=3. Data represent mean ± SEM. Statistical analysis was performed using one-way ANOVA followed by post hoc Tukey’s test (B, F and H) or Student’s unpaired t test (D and J). *P <0.05, **P < 0.01, ***P < 0.001, ****P < 0.0001. ns: not significant. The insets are higher magnification images of the black and yellow boxes. Scale bars: 200 μm (A, E, and G).

### High phosphate-induced intracellular signaling pathways in neutrophil activation

We performed proteomic analysis to identify signaling pathways in neutrophils activated by high phosphate. Lysates of neutrophils exposed to normal (P 0.9mM) or high phosphate (P 2.6mM) were analyzed by LC-MS/MS (Figure 3, A and B). Pathway analysis revealed pathways involving programmed cell death (highlighted in green) and NFE2L2-mediated events (highlighted in blue) (Figure 3C). Then, we focused on the NFE2L2 (Nrf2) pathway, a key antioxidant regulator; its inhibition enhances NET formation by failing to suppress ROS-induced histone citrullination (20–22). We therefore examined whether high phosphate induces ROS production in neutrophils. High phosphate increased intracellular ROS levels in neutrophils, and CKD neutrophils exhibited higher ROS production at both normal and high phosphate levels compared with healthy neutrophils (Figure 3D). PAD4 inhibition (Cl-amidine), which blocks histone citrullination, blocked phosphate-induced NET formation (Figure 3, E and F), indicating that high phosphate promotes ROS production, PAD4 activation, and NET formation.

**Figure 3.**
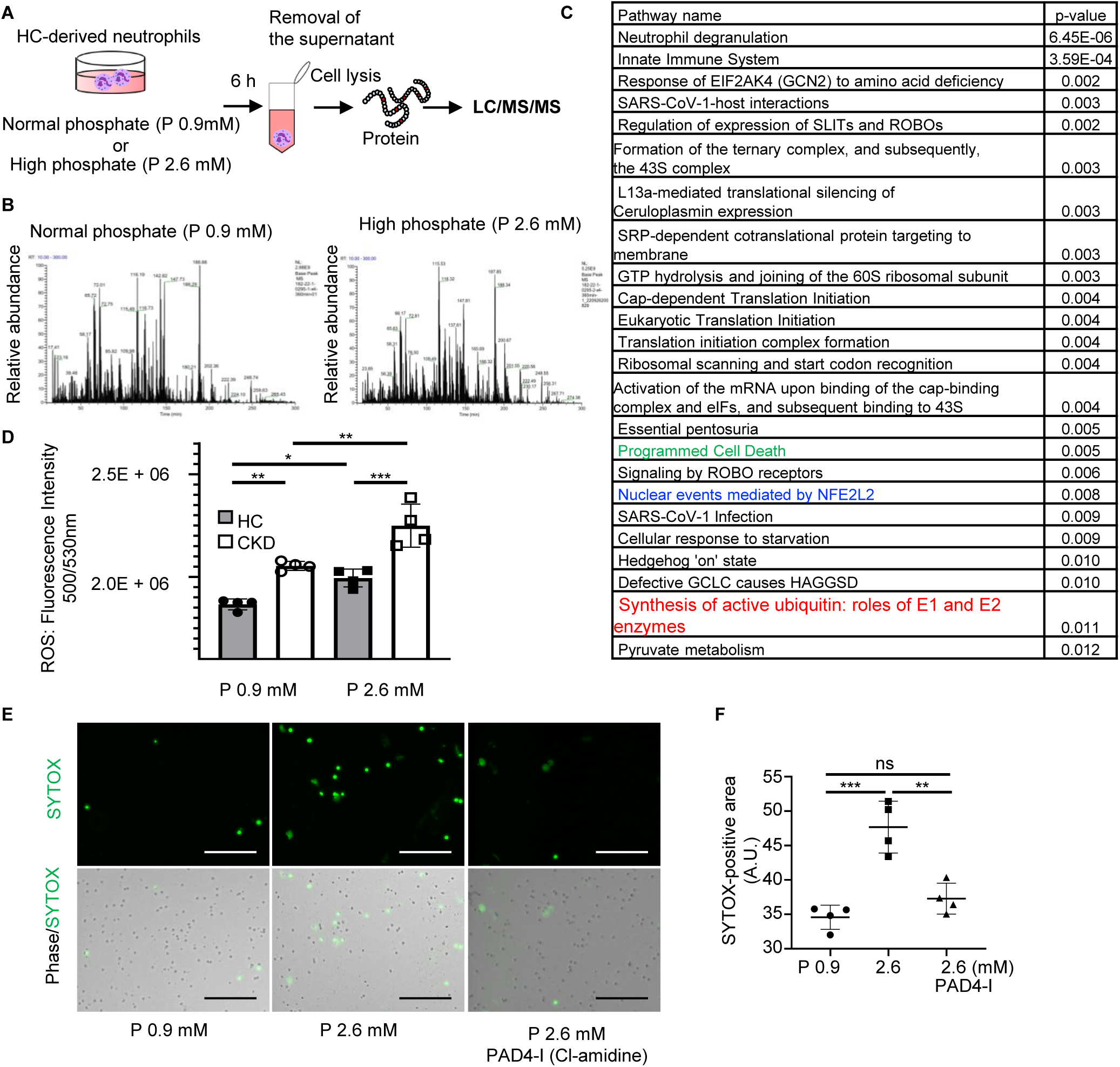
A proteomics analysis of neutrophils in response to high phosphate levels. (A) Neutrophils derived from HC were treated with Normal (P 0.9mM) or high phosphate (P 2.6mM) levels for 6 hours. After centrifugation, the supernatant was removed, and the cells were lysed in lysis buffer. The proteins were digested with trypsin, and each sample was analyzed by LC-MS/MS. (B) The total ion current chromatograms. (C) The most relevant pathways induced by high phosphate, sorted by p-value. (D) The graph quantifies ROS levels. n=4. (E) Neutrophils pretreated with a 200 μM PAD4 inhibitor (PAD4-I, Cl-amidine) were cultured under high phosphate for 6 hours. The panels show representative images of neutrophils stained with SYTOX green. Scale bars: 200 μm (F) The graph quantifies SYTOX-positive area. n=4. Data represent mean ± SEM. Statistical analysis was performed using one-way ANOVA followed by post hoc Tukey’s test (D and F). *P <0.05, **P < 0.01, ***P < 0.001. ns: not significant.

### Fetuin-A depletion by neutrophils in response to high phosphate promotes calcium phosphate crystallization

In this study, we first observed that in the absence of FBS, neutrophils cultured under high phosphate readily formed calcium phosphate crystals, even at relatively low phosphate concentrations (Figure 4A, at phosphate 1.9 mM). To assess the role of FBS, we tested media containing 0–5% FBS and found that crystal formation increased without FBS but was markedly inhibited by FBS, even at high phosphate (Figure 4A). We next evaluated whether neutrophils could promote crystallization in conditions normally resistant to it (2.5% FBS DMEM with P 2.6 mM and Ca 1.8 mM). Remarkably, adding neutrophils induced crystal formation, whereas no crystals formed in their absence (Figure 4B). We then asked whether neutrophil activation under physiological phosphate conditions (2.5% FBS DMEM with P 2.8 mM and Ca 1.8 mM) was driven by phosphate uptake or by micro–calcium phosphate crystals. To address this, we pretreated neutrophils with a NaPi inhibitor, which is an inhibitor of phosphate transport, under FBS-containing conditions. This significantly reduced NET formation, suggesting that phosphate uptake contributes to NET formation rather than secondary crystal deposition (Figure 4C). Given that Fetuin-A in FBS binds calcium and phosphate to inhibit intravascular crystallization (23, 24), we evaluated the change of Fetuin-A levels using HC serum in FBS-free DMEM with P 2.6 mM and Ca 1.8 mM. In the cell (neutrophil)-free conditions, Fetuin-A slightly decreased as phosphate increased, reflecting consumption to prevent crystallization. However, adding neutrophils significantly reduced Fetuin-A levels, especially under high phosphate (Figure 4D). To directly test this, we incubated recombinant Fetuin-A with or without neutrophils in non-crystallizing DMEM (P 0.9 mM, Ca 1.8 mM) for 6 hours. Fetuin-A remained stable in cell-free media but was markedly reduced by neutrophils (Figure 4E). These findings suggest that elevated phosphate levels initiate the reduction of Fetuin-A, a process that is further accelerated by neutrophils. Mechanistically, Fetuin-A has been reported to undergo degradation through ubiquitin–proteasome pathways, including 26S proteasome-associated mechanisms (25). Proteomic analysis revealed robust activation of proteasome-related pathways in neutrophils exposed to high phosphate (Figure 3 C, highlighted in red), with marked upregulation of intracellular 19S regulatory subunits (Rpn2, Rpn3, Rpn6, Rpn9, Rpn12), whereas extracellular 20S components (α7, β5) showed weaker associations (Figure 4F). STRING analysis additionally identified enrichment of proteasome-related molecules, including PSMD8 and PSMD11 (highlighted in red), which are involved in protein degradation pathways (Figure 4G). Collectively, these findings suggest that neutrophils contribute to Fetuin-A depletion, potentially through proteasome-associated pathways, subsequently promoting calcium phosphate crystallization under high phosphate conditions.

**Figure 4.**
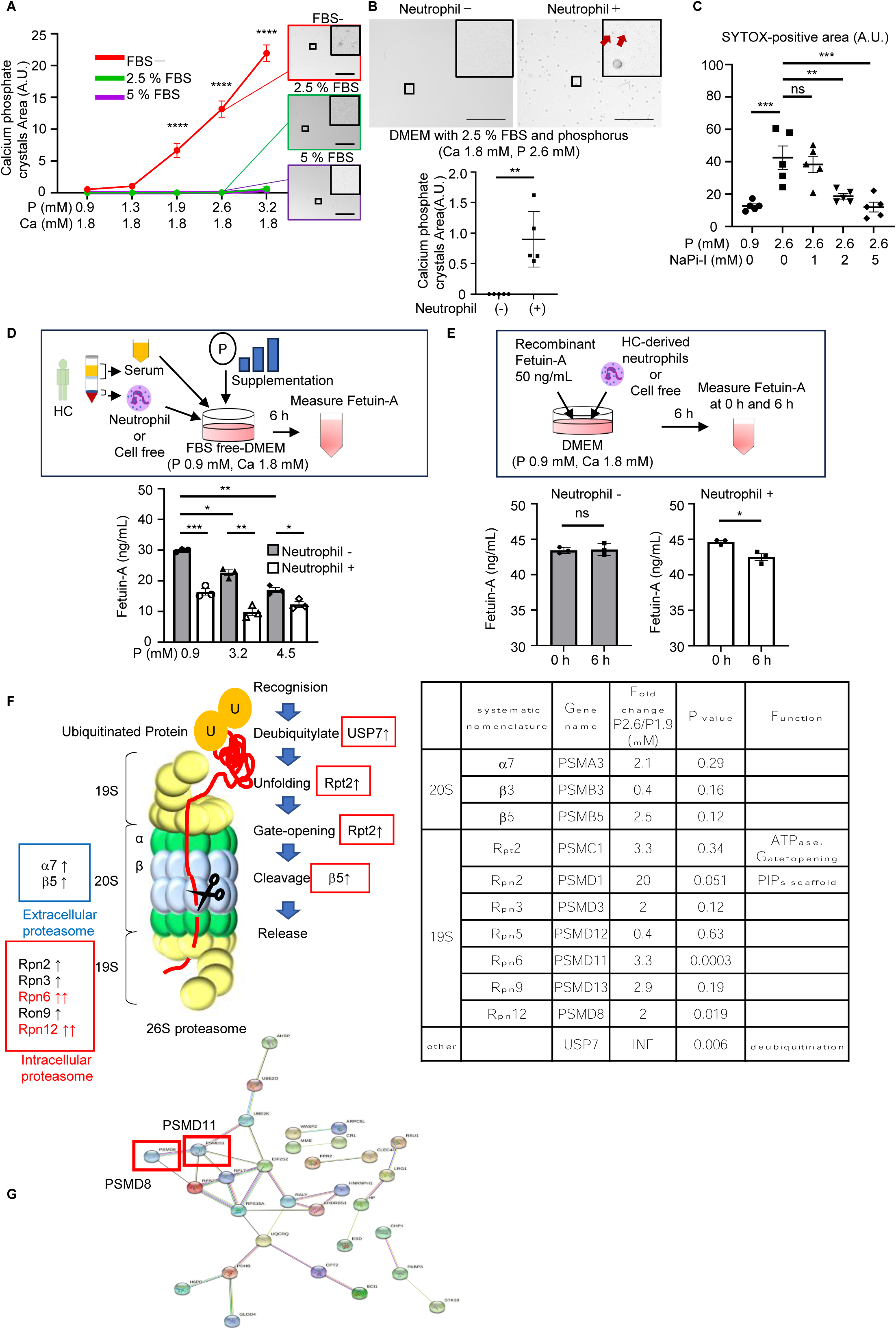
Neutrophil-mediated Fetuin-A depletion and calcium phosphate crystallization. (A) DMEM with various concentrations of phosphate buffer was mixed under conditions without FBS, with 2.5% FBS, and with 5% FBS for 6 hours. The graph shows the quantification of calcium phosphate crystal areas. n=3. (B) DMEM supplemented with 2.5% FBS and phosphate was incubated for 6 hours in the presence or absence of neutrophils. The calcium phosphate crystals are indicated by the red arrows. Below graph shows the quantification of crystal areas. n=5. (C) Neutrophils pretreated with NaPi inhibitor (NaPi-I) were stimulated with high phosphate. Sytox Green fluorescence was quantified after 6 h. (D) DMEM with a variety of phosphate concentrations and serum from HC was cultured with or without healthy neutrophils for 6 hours. Fetuin-A levels in the supernatant were determined by enzyme-linked immunosorbent assay (ELISA). n=3. (E) DMEM supplemented with recombinant Fetuin-A was cultured with and without neutrophils for 6 hours. Fetuin-A levels were quantified at the outset and at the 6-hour mark. n=3. (F) The left image shows a schematic diagram of proteolysis by 26S proteasome and proteins identified through proteomic analysis in neutrophils exposed to high phosphate levels. The intracellular proteasomes and the extracellular proteasomes are indicated by red boxes and blue boxes, respectively. The right table shows the peptides associated with the proteasome, as determined by their fold change in increase and decrease. PIPs: proteasome interacting proteins, INF: infinity. (G) Protein-protein interactions obtained from STRING. Data represent mean ± SEM. Statistical analysis was performed using one-way ANOVA followed by post hoc Tukey’s test (A, C and D) or Student’s unpaired t test (B and E). *P <0.05, **P < 0.01, ***P < 0.001. ns: not significant. The insets are higher magnification images of the black boxes. Scale bars: 200 μm (A and B).

### Calcium phosphate crystals induce massive NET formation

To investigate the effect of crystals on neutrophils, HC neutrophils were incubated with varying phosphate and calcium levels without FBS, a condition that allows crystallization even in cell-free media (Figure 5A). As the concentrations of phosphate and calcium increased, calcium phosphate crystals formed, and neutrophils clustered around these crystals, producing extensive SYTOX- and CitH3-positive NETs (Figure 5, B-E). NET induction was significantly higher in CKD patients’ neutrophils than in HC at all concentrations tested (Supplemental Figure 5, A and B). To determine whether the inhibitory effect of FBS (which contains Fetuin-A) on NET formation and crystallization was due to osmotic pressure, neutrophils were incubated in DMEM supplemented with 2.5% BSA. In BSA-containing media, crystallization and NET formation occurred under high phosphate, unlike in 2.5% FBS (Supplemental Figure 5, C and D), indicating that Fetuin-A, not osmotic effects, mediates this inhibition. To test FGFR1 involvement, neutrophils were treated with an FGFR1 inhibitor under crystallizing conditions. FGFR1 inhibition did not block crystal-induced NET formation (Supplemental Figure 5, E and F), suggesting that NET induction by calcium phosphate crystals occurs independently of phosphate–FGFR1 signaling.

**Figure 5.**
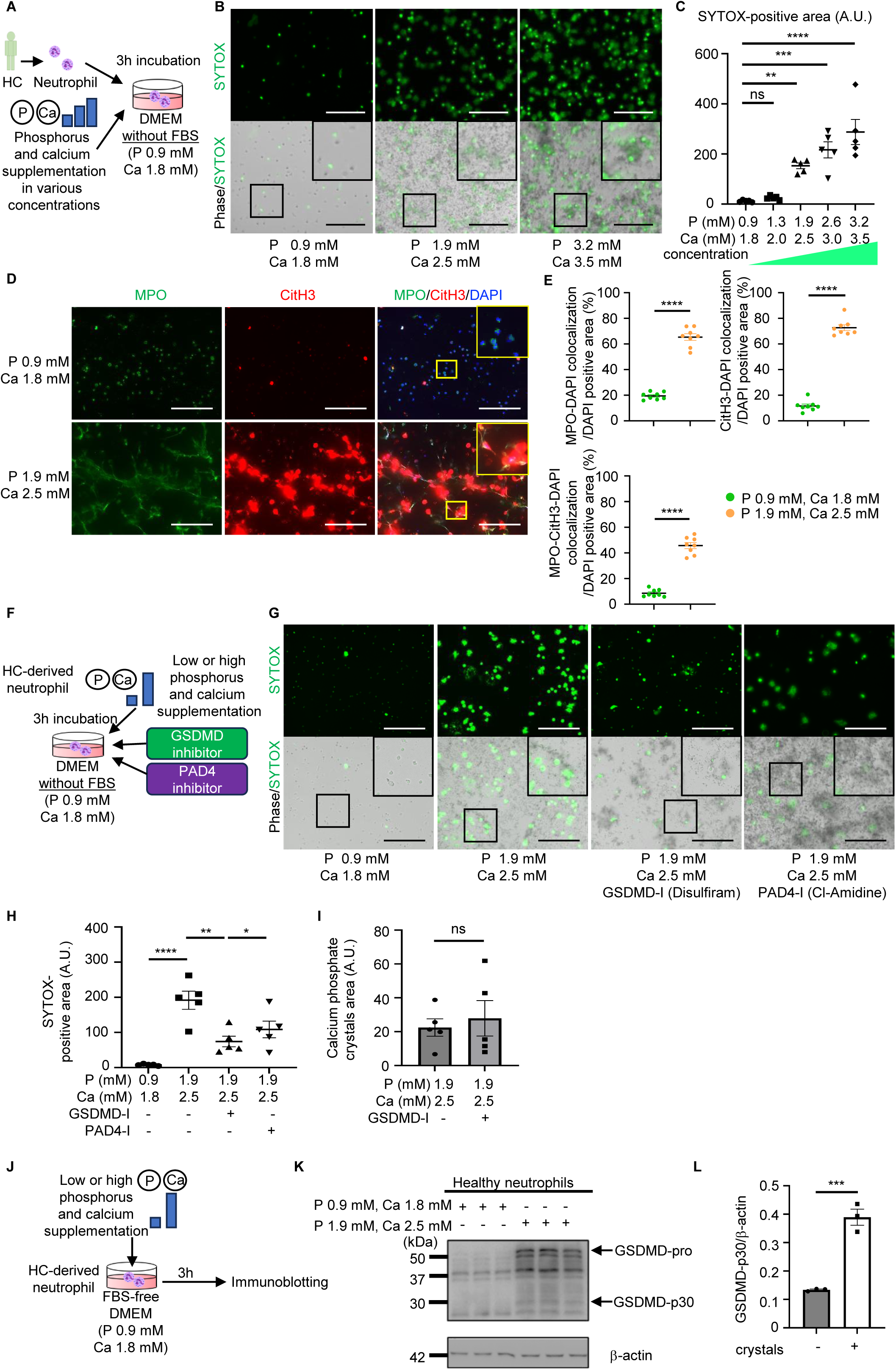
Neutrophil activation in response to calcium phosphate crystals. (A) Neutrophils derived from HC were incubated for 3 hours in DMEM supplemented with various concentrations of phosphate buffer and calcium buffer without FBS. (B) The representative phase-contrast and immunofluorescence images of neutrophils stained by SYTOX green. (C) The quantification of SYTOX-positive area. n=5. (D) The representative immunofluorescence images of MPO (green), CitH3 (red) and DAPI (blue). (E) The quantitative analysis of neutrophils for the MPO-DAPI double-positive area per DAPI positive area, the CitH3-DAPI double-positive area per DAPI positive area, and the MPO-CitH3-DAPI triple-positive area per DAPI positive area, respectively. n=8. (F) Neutrophils pretreated with a 40 μM GSDMD inhibitor (GSDMD-I, Disulfiram) and a 200 μM PAD4 inhibitor (PAD4-I) were incubated for 3 hours in DMEM supplemented with low or high phosphate and calcium buffers without FBS. (G) The representative images of neutrophils. (H) and (I) graphs show the quantification of SYTOX-positive area and calcium phosphate crystal area, respectively (n=5 for each group). (J-L) Neutrophils from HC were incubated with the DMEM supplemented with low or high calcium and phosphate without FBS (J). The protein expression of GSDMD (pro and cleaved p30 (N-terminal fragment)) in neutrophils was detected by immunoblotting using β-actin as a control (K). The graph (L) shows the ratio of GSDMD-p30/β-actin expression. n=3. Data represent mean ± SEM. Statistical analysis was performed using one-way ANOVA followed by post hoc Tukey’s test (C and H) or Student’s unpaired t test (E, I and L). *P <0.05, **P < 0.01, ***P < 0.001, ****P < 0.0001. ns: not significant. The insets are higher magnification images of the black and yellow boxes. Scale bars: 200 μm (B, D, and G).

### Calcium phosphate crystals induce NET formation through the activation of gasdermin D

Proteomic analysis (Figure 3C) indicated that the programmed cell death pathway (highlighted in green) is involved in NET formation induced by high phosphate. Pyroptosis, an inflammatory form of programmed cell death mediated by GSDMD, has been implicated in crystal-induced NET formation (15, 16). We therefore examined GSDMD signaling in response to calcium phosphate crystals. Neutrophils pretreated with a GSDMD inhibitor (Disulfiram) or a PAD4 inhibitor were incubated under conditions promoting crystal formation (high calcium and phosphate without FBS) (Figure 5F). Both inhibitors suppressed NET formation induced by crystals (Figure 5, G and H). Notably, the GSDMD inhibitor had no effect on the extent of crystallization (Figure 5I), indicating that PAD4-GSDMD signaling is involved in calcium phosphate crystal-induced NET formation but not in crystallization itself. Immunoblotting further confirmed increased cleavage of GSDMD in neutrophils stimulated by calcium phosphate crystals (Figure 5, J-L), highlighting the key role of GSDMD in linking crystal formation to NETs.

### High phosphate and calcium phosphate crystal-Induced NETs cause endothelial injury, leading to pro-arteriosclerotic signaling

We elucidated the e□ect of NETs induced by high phosphate and crystals on endothelial injury. In the phosphate-dependent model, neutrophils pretreated with either a FGFR1 inhibitor or a PAD4 inhibitor were incubated in 2.5% FBS DMEM supplemented with low or high phosphate. Crystal-dependent NETs were tested using GSDMD inhibitors in DMEM without FBS. Human umbilical vein endothelial (HuEht) cells exposed to NETs were assessed for damage using calcein-AM/PI staining (Supplemental Figure 6A). Phosphate- and crystal-induced NETs caused endothelial injury. FGFR1 and PAD4 inhibitors protected endothelial cells from phosphate-induced NET injury (Supplemental Figure 6B). GSDMD inhibitors improved endothelial viability in the crystal model (Supplemental Figure 6C). In HuEht cells in the calcium phosphate crystals-dependent model, NETs increased mRNA levels of pro-inflammatory (IL-1β, CXCL1, IL-8) and interferon-inducible proteins (IFIT1). Conversely, the GSDMD inhibitor decreased these expression levels (Supplemental Figure 6D). These findings indicated that NET compositions damage endothelium, leading to the production of inflammatory and arteriosclerotic factors.

### Phosphate-induced neutrophil activation via FGFR1 contributes to intimal calcification in CKD-associated arteriosclerosis

Given our above findings that high phosphate induces NETs via FGFR1, leading to phosphate crystallization, we next investigated whether these phenomena are associated with intimal calcification in human pre-dialysis CKD patients (Supplemental Figure 7A; patient characteristics in Supplemental Table 2) and a murine CKD model. In CKD patients, CD15-positive neutrophils were frequently observed in the intima, whereas few neutrophils were found in non-CKD patients. Additionally, CKD patients showed increased phosphorylation of FGFR1 substrate 2 (pFRS2), a downstream mediator of FGFR1 signaling, and a higher number of CitH3-positive cells within the intima. Quantitatively, CKD patients exhibited more von Kossa-positive calcification and significantly larger pFRS2- and CitH3-positive areas (Supplemental Figure 7, B and C). Immuno-double staining for CD15 and pFRS2 confirmed that pFRS2-positive cells in the intima were predominantly neutrophils (Supplemental Figure 7D). Next, murine CKD vascular calcification model involving intimal and medial lesions, was developed using an adenine (0.25%) and high-phosphate diet, based on a previously described protocol (26). In CKD mice, Ly6b-positive neutrophils are infiltrated in calcified lesions, along with pFRS2 co-localization like human tissues (Supplemental Figure 8). These findings suggest that elevated phosphate levels trigger neutrophil activation via FGFR1 signaling, contributing to intimal calcification through NET formation.

### FGFR inhibition ameliorates intimal vascular calcification in CKD murine model

To evaluate the role of the neutrophil FGFR1 pathway specifically in intimal calcification, we employed a modified model using a lower concentration of dietary adenine (Figure 6A). This model, induced by a diet containing 0.15–0.2% adenine and high phosphate (HP) (see Methods), successfully recapitulates intimal vascular calcification, particularly in the aortic valve. Mice subjected to the modified CKD + HP protocol exhibited elevated serum Cr, BUN, and P levels, compared with control mice receiving a high phosphate diet alone, while serum calcium concentrations remained unchanged. Administration of FGFR inhibitor did not significantly influence these parameters (Figure 6B). Histological analyses revealed substantial interstitial fibrosis and tubular calcium deposition in kidney sections from CKD + HP mice, as demonstrated by Elastica Masson and von Kossa staining, respectively (Figure 6, C and D). These pathological changes were not altered by FGFR inhibitor. Notably, however, intimal calcification of the aortic valve observed in CKD + HP mice was reduced by FGFR inhibition. Furthermore, areas positive for CitH3 tended to be smaller with FGFR blockade (Figure 6, E and F). Collectively, these findings suggest that pharmacological FGFR blockade inhibits NET formation and mitigates intimal vascular calcification in moderate CKD.

**Figure 6.**
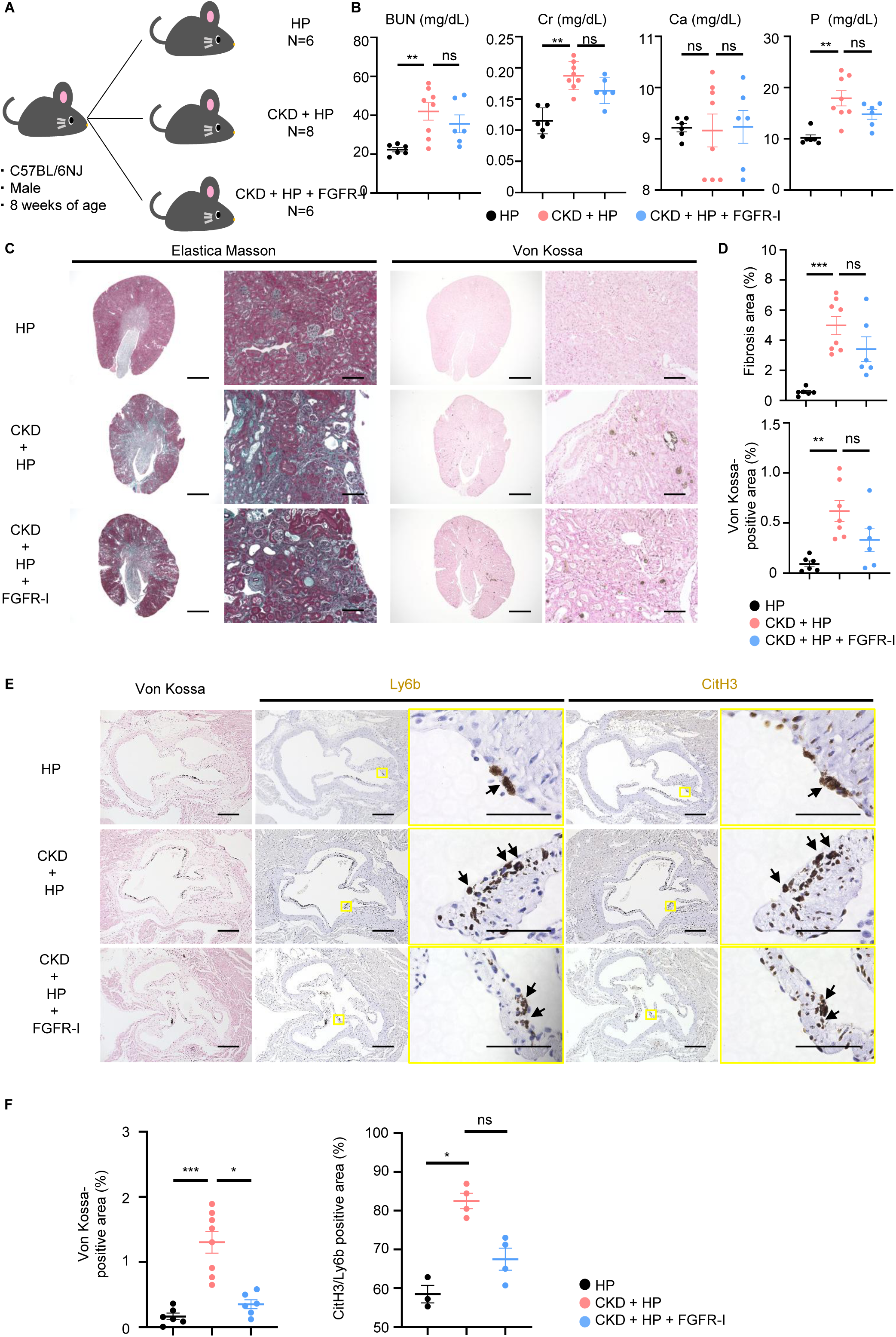
The effects of FGFR inhibitor treatment in CKD-mediated vascular calcification model mice. (A) Illustration of the experimental design. Eight-week-old C57BL/6 male mice were randomly divided into three groups: the high phosphate (HP) group (N=6), the CKD vascular calcification model (CKD + HP) group (N=8) and the FGFR inhibitor-treated CKD vascular calcification model (CKD + HP + FGFR-I) group (N=6). For phosphate loading, mice were administered a high-phosphate diet (1.25% phosphate). CKD was induced by feeding an adenine-rich diet (0.15-0.2% adenine). AZD4547 (FGFR inhibitor, 6mg/kg) or vehicle control was administered via intraperitoneal injection. (B) The serum levels of creatinine (Cr), blood urea nitrogen (BUN), calcium (Ca) and phosphate (P) were measured at the end of the experimental period. (C) The panels show the representative images of Elastica Masson staining and von Kossa staining of kidneys. Scale bars: 1000 μm (low magnification) and 100 μm (high magnification). (D) The graphs show the percentage of fibrosis area and von Kossa-positive area per section. (E) The representative images of von Kossa, Ly6b (brown), and CitH3 (brown) staining of the aortic valve. The black arrows indicate Ly6b- and CitH3-positive cells, respectively. The insets are higher magnification images of the yellow boxes. Scale bars: 200 μm (low magnification) and 50 μm (high magnification). (F) The graphs show the percentage of von Kossa-positive area calcified area in the aortic valve annulus (HP: n=6, CKD + HP: n=8, CKD + HP + FGFR-I: n=6) and the ratio of CitH3-positive area to Ly6b-positive area within the aortic valve, reflecting NET-forming neutrophils (HP: n=3, CKD + HP: n=4, CKD + HP + FGFR-I: n=4). Data represent mean ± SEM. Statistical analysis was performed using Kruskal-Wallis test (B, D and F). *P <0.05, **P < 0.01, ***P < 0.001. ns: not significant.

### Neutrophils respond to the efflux of phosphate from necrotic tissue

We hypothesized that neutrophils respond to phosphate released from necrotic cells as an alarmin. In vitro, necrotic endothelial cells released significantly higher levels of phosphate compared to controls (Supplemental Figure 9). These findings suggest that phosphate released from necrotic tissue may contribute to neutrophil activation and subsequent NET formation.

## Discussion

In this study, we identified neutrophil activation via FGFR1 as a key pathway linking phosphate excess to NET formation and intimal vascular calcification. Our findings reveal that neutrophils expressing FGFR1 respond to elevated phosphate levels by forming NETs via ERK phosphorylation, ROS production, and PAD4 activation. During these steps, neutrophils deplete Fetuin-A through proteasomal activity, promoting calcium-phosphate crystallization. Subsequently formed crystals induce a secondary wave of NET formation through GSDMD signaling, exacerbating vascular damage and promoting pro-inflammatory and pro-arteriosclerotic pathways. Notably, neutrophils from CKD patients exhibited heightened ROS production and sensitivity to phosphate compared to those from healthy controls, leading to more robust NET formation. This enhanced responsiveness may reflect chronic exposure to systemic inflammation and hyperphosphatemia in CKD. These results suggest a pathological feedback loop wherein calcification and NET formation perpetuate each other, driving vascular calcification in CKD. In line with previous reports showing that CKD patients have a higher basal capacity to form NETs (27, 28), our study provides functional evidence that phosphate directly triggers NET formation in CKD neutrophils. Furthermore, recent studies have demonstrated that osteocytes sense phosphate via FGFR1 and regulate phosphate homeostasis via FGF23 secretion (10, 29). Extending this concept to innate immunity, we demonstrate that high phosphate activates neutrophil FGFR1, initiating ERK signaling and NET formation. Moreover, analysis of calcified arteriosclerotic lesions from both human CKD patients and vascular calcification model mice revealed the infiltration of neutrophils exhibiting activated FGFR1 signaling and the formation of NETs within calcified vascular lesions. Although FGFR1 contributes to phosphate-induced NET formation, its relationship with NaPi-mediated phosphate transport remains unresolved. NaPi inhibition also attenuated NET formation, suggesting cooperative signaling. Although the proportion of neutrophils in the vascular tissues was relatively low, this may reflect the chronic nature of arteriosclerosis in CKD (28), where persistently elevated circulating neutrophils could exert cumulative effects on the vessel wall over time.

A surprising discovery in our study was that simply adding neutrophils to the medium significantly promoted calcium-phosphate precipitation. Fetuin-A, a protein that prevents calcium-phosphate crystallization by binding calcium and phosphate (30), was found to be decreased by neutrophils. Proteomic analysis revealed that high phosphate levels activate the neutrophil intracellular proteasome, which may facilitate the degradation of extracellular Fetuin-A internalized by neutrophils, thereby promoting crystal formation. Additionally, we demonstrated that calcium-phosphate crystals induce massive NET formation in neutrophils via GSDMD/PAD4-mediated signaling, which is distinct from the phosphate-FGFR1 pathway (see graphic abstract). Both signaling pathways may serve as potential therapeutic targets for mitigating vascular calcification and inflammation in CKD. Importantly, our in vivo study demonstrated that FGFR1 inhibition substantially attenuated NET formation and intimal vascular calcification in a murine CKD model, accompanied by suppression of FGFR1 signaling. These findings highlight phosphate-sensing neutrophils as a potential therapeutic target for CKD-associated vascular calcification, for which effective therapies remain unavailable.

Finally, to explain why neutrophils (immune cells) respond to phosphate, we hypothesized that phosphate released from necrotic tissue serves as an immune danger signal, triggering NET formation to sequester pathogens. In vitro, necrotic endothelial cells released higher levels of phosphate compared with controls. Such phosphate release could serve as a trigger for neutrophil activation and subsequent calcification. Neutrophils are known to infiltrate necrotic tissues at an early stage (31), supporting the notion that they respond to phosphate and promote calcification through NET formation. Chronic inflammation has long been implicated in vascular calcification (32), and previous studies demonstrated that neutrophils form NETs and promote calcification in necrotic lesions such as tuberculosis, effectively sequestering pathogens (33, 34). These observations raise the possibility that neutrophils sense phosphate as an alarmin and contribute to pathological calcification in CKD, analogous to how immune thrombosis contains injury in other settings (35). However, in CKD patients, persistent elevation of serum phosphate exacerbates this immune-calcification process through persistent neutrophil activation and dysregulated phosphate metabolism.

Several limitations should be acknowledged. Although this study focused on neutrophils, histological analysis revealed only modest neutrophil infiltration in calcified lesions, which may reflect the chronic nature of arteriosclerosis in CKD. In addition, while we used pharmacological FGFR inhibition to evaluate the role of neutrophil phosphate sensing, FGFR1 is also expressed in other cell types, and off-target effects cannot be excluded. Previous reports have described that FGFR inhibition can increase serum phosphate levels (29), but in our CKD mouse model phosphate elevation was not observed. One possible explanation is that FGFR inhibition may have exerted nephroprotective effects, leading to enhanced phosphate excretion, possibly through attenuation of renal injury. Finally, in moderate CKD patients, where serum phosphate levels are often not overtly elevated, vascular calcification may still progress. Prior studies have reported decreased fetuin-A concentrations in CKD (30), which, in combination with mild phosphate elevations, could create a permissive environment for local calcium–phosphate deposition.

In conclusion, this study identifies neutrophil activation via FGFR1 and GSDMD as critical drivers of vascular calcification in CKD. By demonstrating the involvement of neutrophils in immune-mediated calcification, we provide a novel perspective on the immunological contributions to CKD-associated vascular pathology. These findings may inform the development of innovative therapies to address the cardiovascular complications of CKD.

## Methods

### Sex as a biological variable

We examined both male and female human neutrophils. Findings were similar for both sexes. Our study examined male mice because male animals exhibited less variability in phenotype.

### Patients and Blood Samples

Patients enrolled in the experiment on the addition of high levels of phosphate to neutrophils included four non-dialysis CKD patients and four healthy controls (HC) (Supplemental Table1). For CKD and healthy control comparison experiments, neutrophils from four CKD patients or four healthy controls were processed independently. For stimulation and inhibition studies, neutrophils isolated from a single donor were seeded into 4–8 wells per condition to assess technical reproducibility, and the same experiments were independently repeated using neutrophils from multiple donors to confirm reproducibility across biological replicates. Peripheral blood was collected in heparinized tubes. The creatinine (Cr), blood urea nitrogen (BUN), Albumin, HbA1c, calcium (Ca), and phosphate (P) level were measured using the JCA-BM8040 (Japan Electron Optics Laboratory).

### Neutrophil isolation and NET induction

Neutrophils were isolated from the samples using density-gradient centrifugation with Polymorphprep (Axis-Shield). Subsequently, the cells were seeded onto a 96-well plate (1 × 10^5^ cells/ml) and exposed to varying stimuli (4–8 wells per condition). Stimulation with phosphate and calcium was performed using 0.2 M phosphate buffer (pH 7.3) containing Na_2_HPO_4_ and NaH_2_PO_4_, and 0.1 M calcium chloride buffer. For experiments involving high phosphate, neutrophils were incubated with 2.5% fetal bovine serum (FBS) DMEM (Sigma-Aldrich) media (containing baseline 0.9 mM phosphate and 1.8 mM calcium) with varying phosphate concentrations (0.9–3.2 mM phosphate) for 6 hours. For calcium phosphate crystal induction assay, calcium and phosphate solutions were added to DMEM without FBS. The concentrations of phosphate and calcium were adjusted to final concentrations of 3.2 mM phosphate and 2.5 mM calcium. Neutrophils were cultured in the media for 3 hours at 37℃ in 5% CO₂ atmosphere. Under these FBS-free conditions, spontaneous crystal formation was observed by phase contrast microscopy. NET formation was assessed using SYTOX Green (Thermo Fisher) fluorescence imaging and immunocytochemistry for myeloperoxidase (MPO; Origine) and citrullinated histone H3 (CitH3; Abcam) positivity. For ROS production, neutrophils were stained with DCFH-DA (Dojindo) under various phosphate levels. For inhibitory experiments, neutrophils were pretreated with the FGFR1 inhibitor SU5402 (20 µM; Sigma-Aldrich), the GSDMD inhibitor Disulfiram (40 µM; MedChemExpress), the PAD4 inhibitor Cl-amidine (200 µM; Sigma-Aldrich), or the NaPi inhibitor (Phosphonoformic acid, concentrations of 1–5 mM) for 0.5 or 1 hour prior to incubation in high phosphate or calcium phosphate crystal-forming conditions. The quantification of positive area was performed using ImageJ/Fiji software (NIH). Multiple fields of view were obtained for each well, and the mean value was calculated. The area of calcium phosphate crystals was quantified by ImageJ software from phase contrast images, excluding areas occupied by neutrophils. Image analysts were blinded to group allocation.

### Isolation and stimulation of human monocyte and lymphocyte

Peripheral Blood Mononuclear cells (PBMCs) were isolated from healthy human donors by density centrifugation using Ficoll-Paque Plus (GE Healthcare). Monocytes were isolated from PBMCs using CD14 MicroBeads (Miltenyi Biotech) and magnetic columns, according to the manufacturer’s instructions. The cells obtained through negative selection were identified as lymphocytes. Lymphocytes and monocytes were seeded into 96-well plates at a density of 1 × 10^5^/mL, respectively, and then incubated for 6 hours in 2.5% FBS DMEM medium with various concentrations of phosphate buffer. Cells were stained with SYTOX Green.

### In vitro immunocytochemistry

Cells were fixed with 4% paraformaldehyde for 15 minutes at room temperature, permeabilized with 0.2% Triton X-100 for 10 minutes, and blocked with 5% BSA. Cells were incubated overnight at 4°C with primary antibodies against MPO (Origene) and CitH3 (Abcam), followed by appropriate Alexa Fluor-conjugated secondary antibodies. Nuclear counterstaining was performed with DAPI (Vector Laboratories). Fluorescence images were acquired using a fluorescence microscope (Keyence).

### FGFR1 Knockdown in HL-60 Cells Using Small Interfering RNA (siRNA)

Small interfering RNA (siRNA) targeting FGFR1 and non-targeting control siRNA were purchased from Thermo Fisher Scientific (Silencer Select siRNA, USA). HL-60 cells were induced to neutrophilic differentiation by treatment with 1 μM all-trans-retinoic acid (ATRA) for 24 hours. The cells (1 × 10^6^ cells in 10 μL Neon tip) were resuspended with 500 nM of FGFR1 siRNA or non-targeting control siRNA, and transfected using the Neon Transfection System (Thermo Fisher Scientific) under the following optimized electroporation parameters: a single pulse at 1350 V for 35 ms, according to the manufacturer’s instructions for HL-60 suspension cells. The target sequence: CUCACUGUGGAGUAUCCAU, AUGGAACUCCACAGUGAG.

Transfections were performed twice, at 24 h and 48 h after the initiation of ATRA treatment. At 72 h (24 h after the second transfection), cells were harvested for quantitative real-time PCR to confirm FGFR1 knockdown efficiency, which typically ranged from −60% to −85%. At 24 hours after the second transfection, the cells were treated with high phosphate (2.6 mM) or control conditions, and then incubated for a period of 10 hours. NET formation was assessed by SYTOX Green fluorescence and immunocytochemistry for CitH3. The cell viability was confirmed by trypan blue (Thermo Fisher Scientific).

### Quantitative real-time PCR

Total RNA was extracted using TRIZOL reagent (Thermo Fisher Scientific) according to the manufacturer’s recommendations. RNA was reverse transcribed into first-strand cDNA using SuperScript III (Thermo Fisher Scientific). Quantitative real-time PCR was performed using Fast SYBR Green Master Mix (Thermo Fisher Scientific). Gene expression values were normalized to GAPDH as a housekeeping gene. FGFR1 primers were used for HL-60 cells, whereas IL-1β, IFIT1, CXCL1, and IL-8 primers were used for HuEht cells. The primers used were as follows:

GAPDH F: GGGAAGCTTGTCATCAATGGA

R: TCTGGCTCCTGGAAGATGGT
FGFR1 F: CGCCCCTGTACCTGGAGATCATCA

R: TTGGTACCACTCTTCATCTT
IL-1β F: CCACAGACCTTCCAGGAGAATG

R: GTGCAGTTCAGTGATCGTACAGG
IFIT1 F: AAAAGCCCACATTTGAGGTG

R: GAAATTCCTGAAACCGACCA
CXCL1 F: AGGGAATTCACCCCAAGAAC

R: ACTATGGGGGATGCAGGATT
IL-8 F: GAGAGTGATTGAGAGTGGACCAC

R: CACAACCCTCTGCACCCAGTTT

### Immunoblotting analysis

Treated neutrophils or HL-60 cells were homogenized and lysed in buffer containing 20 mM Tris-HCl, 150 mM NaCl, 1% Triton, 1 mM EDTA, 1X protein inhibitor cocktail (Roche Life Science), and phosphatase inhibitor cocktail (Sigma-Aldrich). The proteins were separated by SDS-PAGE and transferred to polyvinylidene fluoride membranes. Membranes were incubated with 5% skimmed milk, probed with primary antibodies against GSDMD (Cell Signaling Technology), ERK (Cell Signaling Technology), phospho-ERK (Cell Signaling Technology), and β-actin (Cell Signaling Technology) overnight at 4°C, and incubated with horseradish peroxidase (HRP)-conjugated secondary antibodies (Thermo Scientific). Protein signals were detected using chemiluminescence (GE Healthcare), and densitometric analysis was conducted with ImageJ software.

### Proteomics analysis of neutrophils

Neutrophils derived from healthy controls were treated with phosphate concentrations of 0.9 mM or 2.6 mM for 6 hours. Following centrifugation, the supernatant was removed, and cells were lysed in lysis buffer. Proteins were digested with trypsin and analyzed by LC-MS/MS. Protein-protein interactions were examined using STRING database analysis.

### Measurement of extracellular Fetuin-A levels

A variety of phosphate concentrations and serum from a healthy control were added to DMEM. The medium was cultured in the presence or absence of neutrophils for 6 hours. Fetuin-A levels in the supernatant were determined by enzyme-linked immunosorbent assay (ELISA; BioVendor). The concentration of recombinant Fetuin-A was measured before and after 6 hours of incubation in DMEM with or without neutrophils.

### Endothelial cell injury assay

Human umbilical vein endothelial cells (HuEht) were cultured in HuMedia-EG2 (Kurabo) and seeded onto the bottom layer of 24-well plates. After overnight culture, HuMedia-EG2 medium was replaced with DMEM containing 2.5% FBS or FBS-free. To avoid direct interactions with neutrophils, neutrophils exposed to high phosphate or calcium-phosphate crystals were added to transwell inserts with a 0.4 µm pore size, which allowed the diffusion of soluble factors but prevented neutrophil infiltration. After 3 or 6 hours of co-incubation, endothelial cell damage caused by neutrophil-derived factors was assessed using calcein-AM/propidium iodide staining (Dojindo) and quantified using fluorescence microscopy. The level of cell toxicity was determined by calculating the percentage of calcein-positive cells out of the total sum of calcein- and PI-positive cells.

### Analysis in human tissues

Formalin-fixed, paraffin-embedded Thoracic and abdominal aorta tissues from CKD and non-CKD patients (eGFR ≥60 mL/min/1.73 m² and no albuminuria by record review) with arteriosclerosis undergoing graft replacement were sectioned for histological analysis. HE and von Kossa staining were performed to assess calcification. For immunohistochemistry, sections were deparaffinized, rehydrated, and subjected to antigen retrieval in sodium citrate buffer (pH 6.0). Endogenous peroxidase activity was blocked with 3% H2O2, followed by blocking with 10% goat serum at room temperature for 1 hour. Sections were incubated overnight at 4°C with anti-CitH3 antibody (abcam) and anti-pFRS2 antibody (R&D Systems), followed by secondary antibody and Histofine DAB substrate (Nichirei). For the double staining, CD15 (Dako) and pFRS2 were used as primary antibodies, followed by Histofine DAB substrate and Fuchsin substrate-chromogen systems (Dako) reacted with alkaline phosphatase polymer (Dako). Positive areas were quantified using ImageJ/Fiji software.

### Animal studies

C57BL/6 mice were maintained under specific pathogen-free conditions, and the experiments described below conformed to the Hokkaido University Guidelines for the Care and Use of Laboratory Animals. Mice were purchased from the National Institute of Biomedical Innovation, Health, and Nutrition (Osaka, Japan). Experimental checklist is based on the ARRIVE guidelines 2.0. The evaluators performing the assessments were kept unaware of the group allocations. Blinding was introduced as a measure to mitigate potential observer bias in the recorded outcome measurements. Mice were also excluded if they died prematurely, preventing the collection of accurate biochemical and histological data. These criteria were established a priori. Serum levels of P, Ca, Cr, and BUN were determined by Oriental Yeast Co., LTD. Nagahama LSL. The animal experiments were approved by the Hokkaido University Animal Experiment Committee.

### High-adenine pilot model for CKD-associated intimal and medial calcification (qualitative histology only)

Eight-week-old male C57BL/6J mice were fed a high-phosphate diet (1.25% phosphate) for 13 weeks together with an intermittently administered adenine diet (0.25% adenine) over the same period, as previously described (26). This protocol effectively induced both intimal and medial vascular calcification, but was associated with high mortality; therefore, data from this model are presented solely as qualitative histology and were not used for statistical inference. Aortae were processed as consecutive 5-μm paraffin sections. Vascular calcification was assessed using von Kossa staining on aortic sections. Neutrophil infiltration and FGFR1 activation were evaluated by immunohistochemistry for Ly6B (Bio-Rad) and phospho-FRS2 (R&D Systems).

### Administration of a FGFR inhibitor in a CKD intimal vascular calcification mouse model

To evaluate the role of the neutrophil FGFR1 pathway specifically in intimal calcification, we employed a modified model using a lower concentration of dietary adenine. Eight-week-old male C57BL/6 mice were fed a high-phosphate diet (1.25% phosphate) for 13 weeks, in combination with an intermittently adenine-containing diet (0.15%-0.2%) over the same period as previously described (36, 37). In this model, vascular calcification is not observed in the aortic wall but is confined to the aortic valve annulus, thereby allowing selective assessment of intimal lesions. Mice were randomly allocated to the three groups: high-phosphate diet alone (HP, n=6), CKD intimal vascular calcification model (CKD + HP, n=8), and FGFR inhibitor-treated CKD intimal vascular calcification model (CKD + HP + FGFR-I, n=6). FGFR inhibitor (AZD4547, 6 mg/kg; MedChemExpress) (38) or vehicle was administered via intraperitoneal injection beginning at 15 weeks of age. FGFR inhibitor treatments were given three times per week until the mice reached 21 weeks of age as previously described. Kidney injury was assessed based on histological findings, using von Kossa staining and Elastica Masson staining. Additionally, blood urea nitrogen (BUN), creatinine, calcium and phosphate levels were measured at the end of the experimental period. Vascular calcification was assessed using von Kossa staining on aortic valve sections. Neutrophil infiltration and formation of NETs were evaluated by immunohistochemistry for Ly6B (Bio-Rad) and CitH3 antibody (abcam).

### Phosphate concentration changes induced by cell death

HuEht cells treated with 5 mM hydrogen peroxide (H₂O₂) were incubated in DMEM medium for 4 hours to induce cell necrosis. Subsequently, the concentrations of phosphate released from cells in the supernatant were measured.

### Statistics

Data are presented as mean ± SEM. For comparisons between two groups, Student’s t-test was used. Differences between greater than 2 groups were analyzed by 1-way ANOVA followed by Dunnett’s multiple-comparison test, one-way ANOVA with Tukey’s post hoc test and Kruskal-Wallis test. A p-value < 0.05 was considered statistically significant. Statistical analysis was performed using GraphPad Prism version 9.5.0 for Windows (GraphPad Software).

### Study approval

Studies involving human materials were approved by the Hokkaido University Hospital Ethics Committee (approval nos. 021-0057 and 024-0071), and written informed consent was obtained from all participants. Animal experiments were approved by the Hokkaido University Animal Experiment Committee (approval no. 25-0071).

### Data availability

In line with our commitment to transparency and the advancement of scientific knowledge, we provide access to relevant data to support the findings presented in this study. The complete dataset, including raw data and analysis scripts, is available as Supplemental material for this publication. For inquiries regarding data access or requests for additional information, interested researchers are invited to contact the corresponding author at.

## Supporting information

graphic abstract

## Author contributions

A.M.H. and D.N. performed most experiments and wrote the manuscript. A.M.H., M.K., T.K., and S.T. analyzed data. T.S., K.W.-K., F.H., S.N., S.S.-A., Y.U., M.K., S.M., Y.N., U.T., Y.S., S.W., A.I., and T.A. contributed to data acquisition, interpretation, or provided critical input. D.N. A.I., and T.A. supervised the study. All authors read and approved the final manuscript.

## Acknowledgments

We thank the staff of the Institute for Animal Facility, Faculty of Medicine and Graduate School of Medicine, Hokkaido University. We also thank the staff from Editage Group for editing the draft of this manuscript. This study was supported by a Grant-in-Aid from the Ministry of Education, Culture, Sports, Science, and Technology of Japan (grant number JP 20 24K11423 and JP 20K08581).

## Notes

### Competing Interest Statement

The authors have declared no competing interest.

