## Supplementary figures and images for "FGFR1-phosphate sensing and crystal-induced gasdermin D signaling in neutrophils drive vascular calcification in CKD"

### graphic abstract

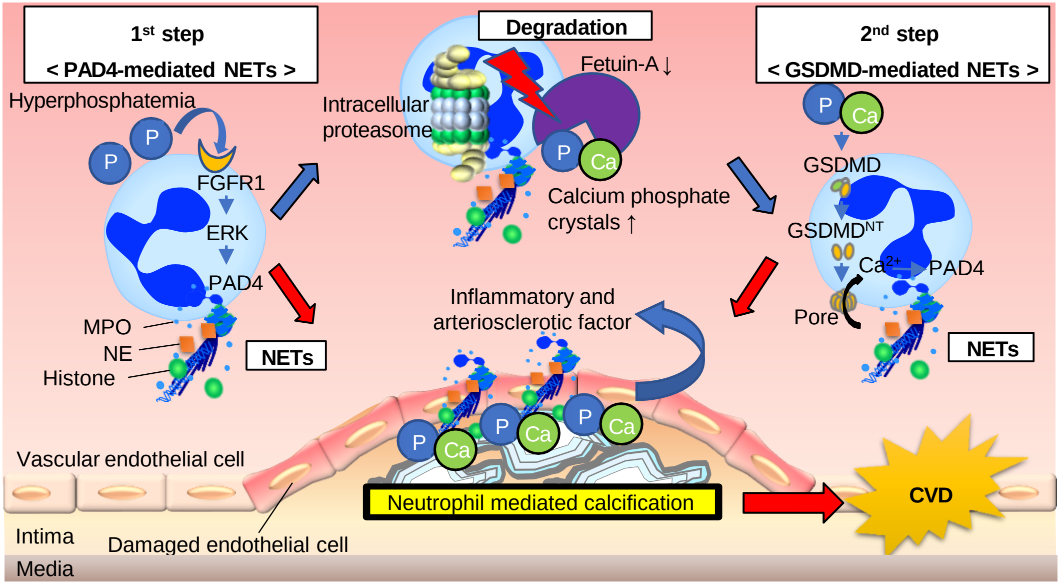
