## Supplemental figure for "FGFR1-phosphate sensing and crystal-induced gasdermin D signaling in neutrophils drive vascular calcification in CKD"

Supplemental Figure 1

**A**

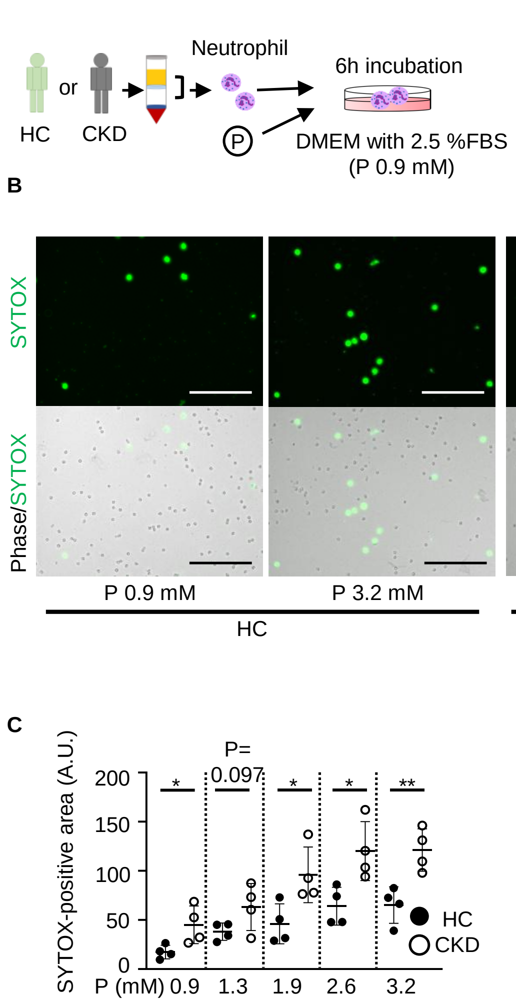

Supplemental Figure 2

A

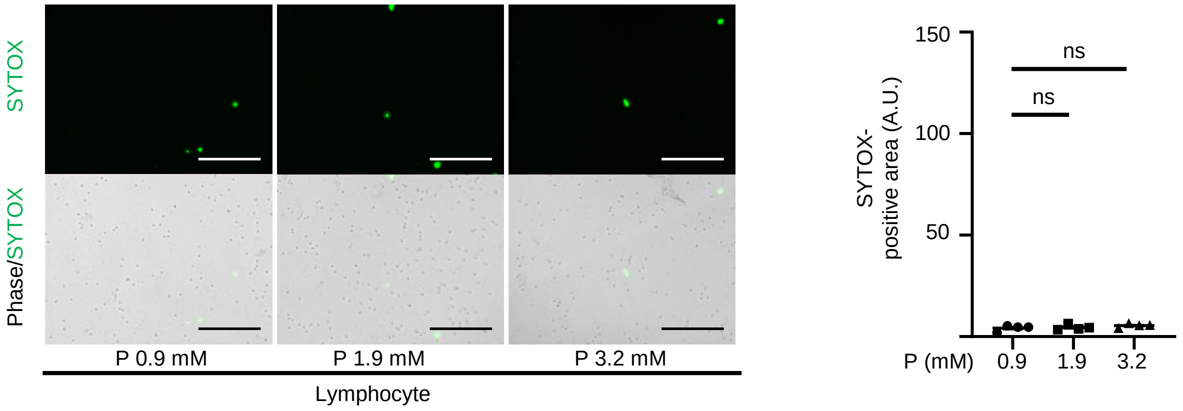


B

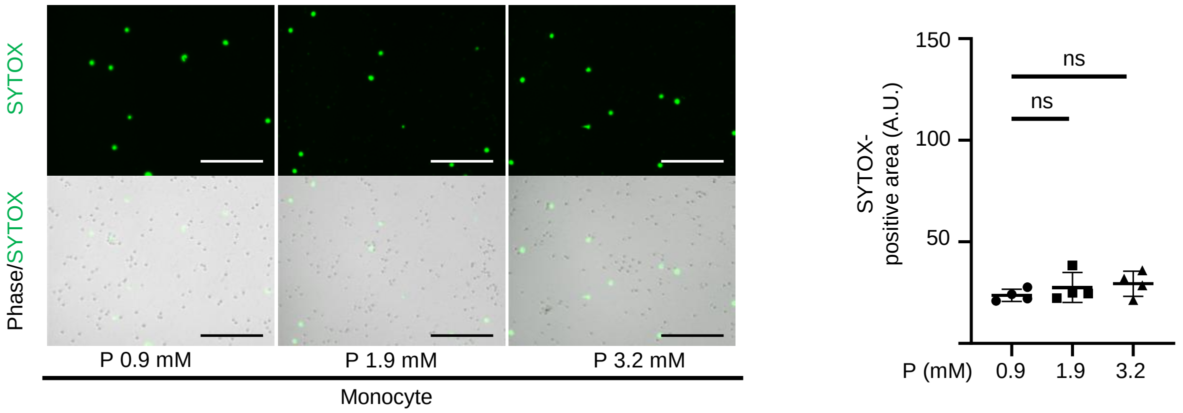

Supplemental Figure 3

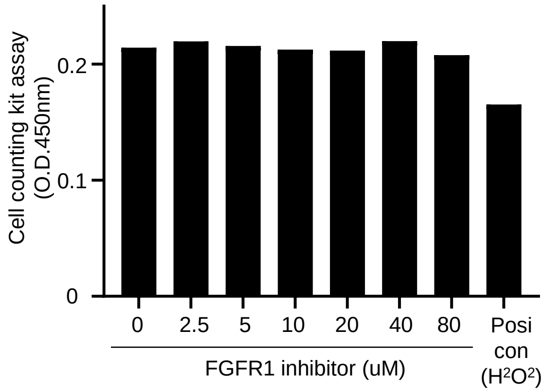

Supplemental Figure 4.

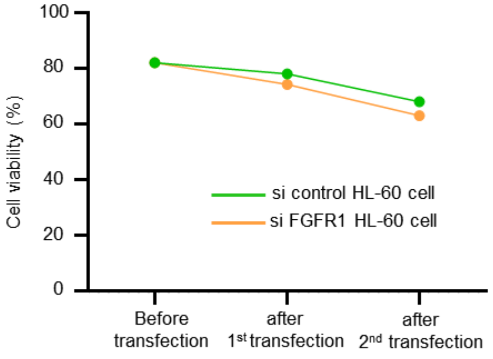

Supplemental Figure 5

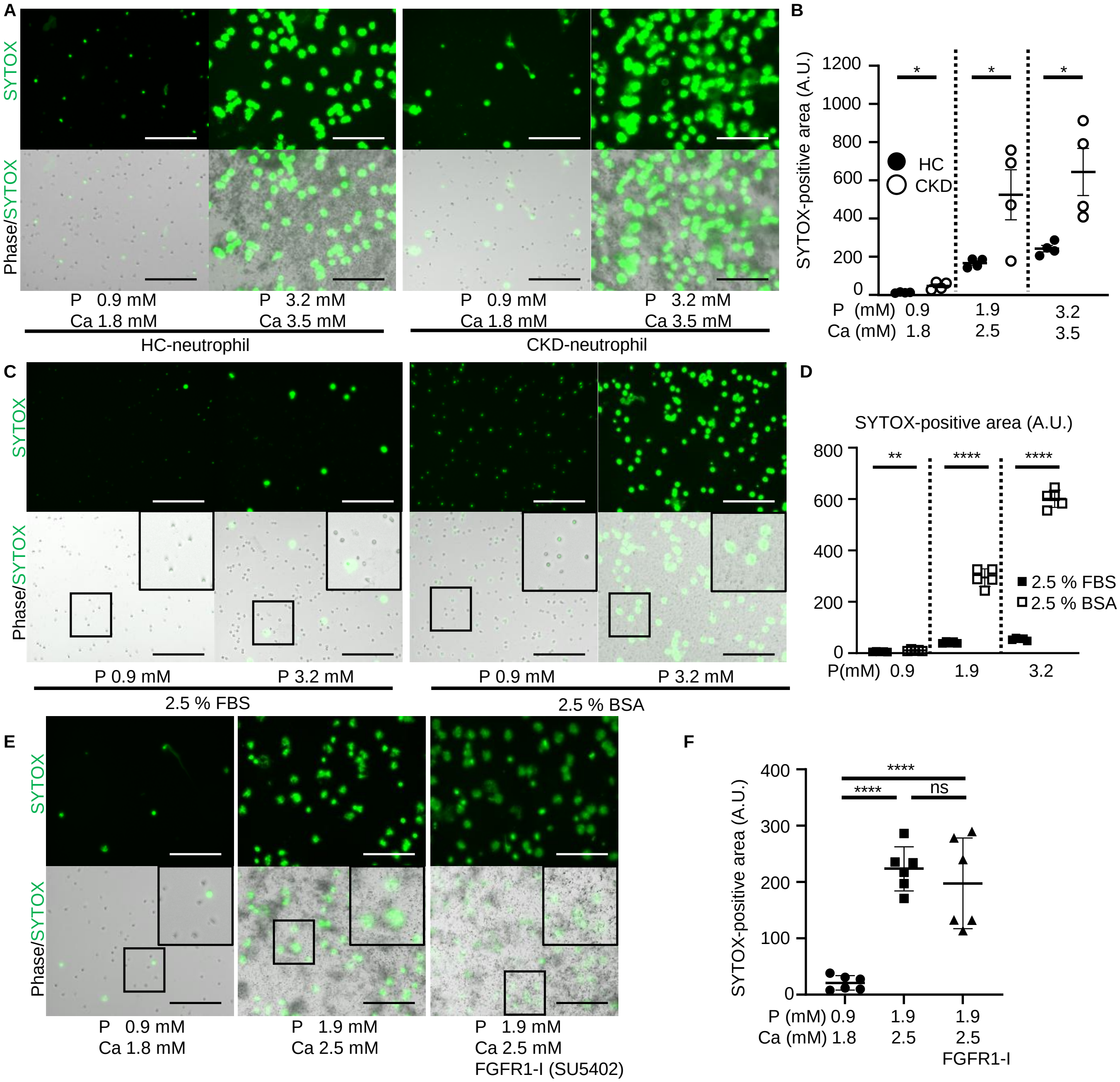

Supplemental Figure 6

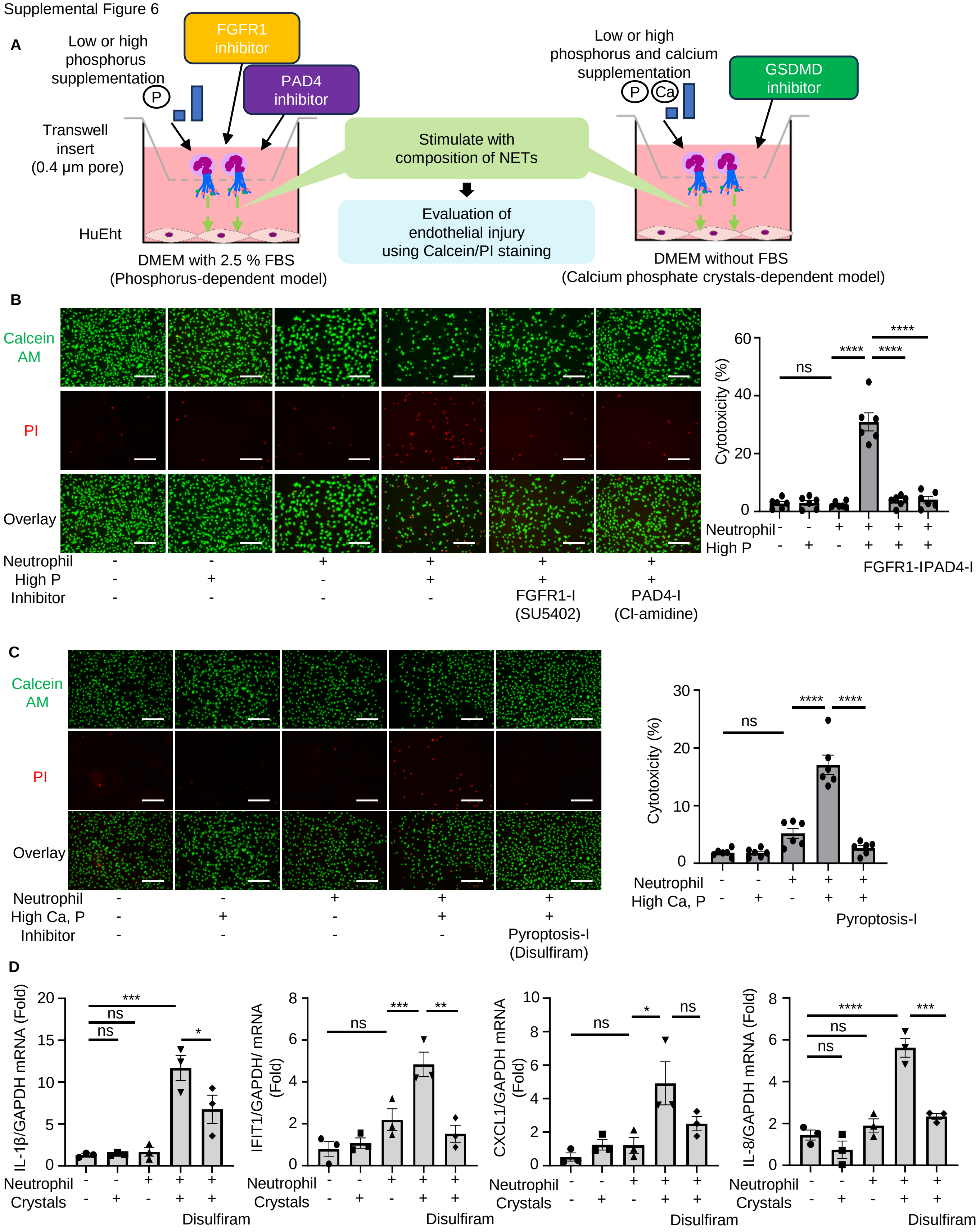

Supplemental figure 7

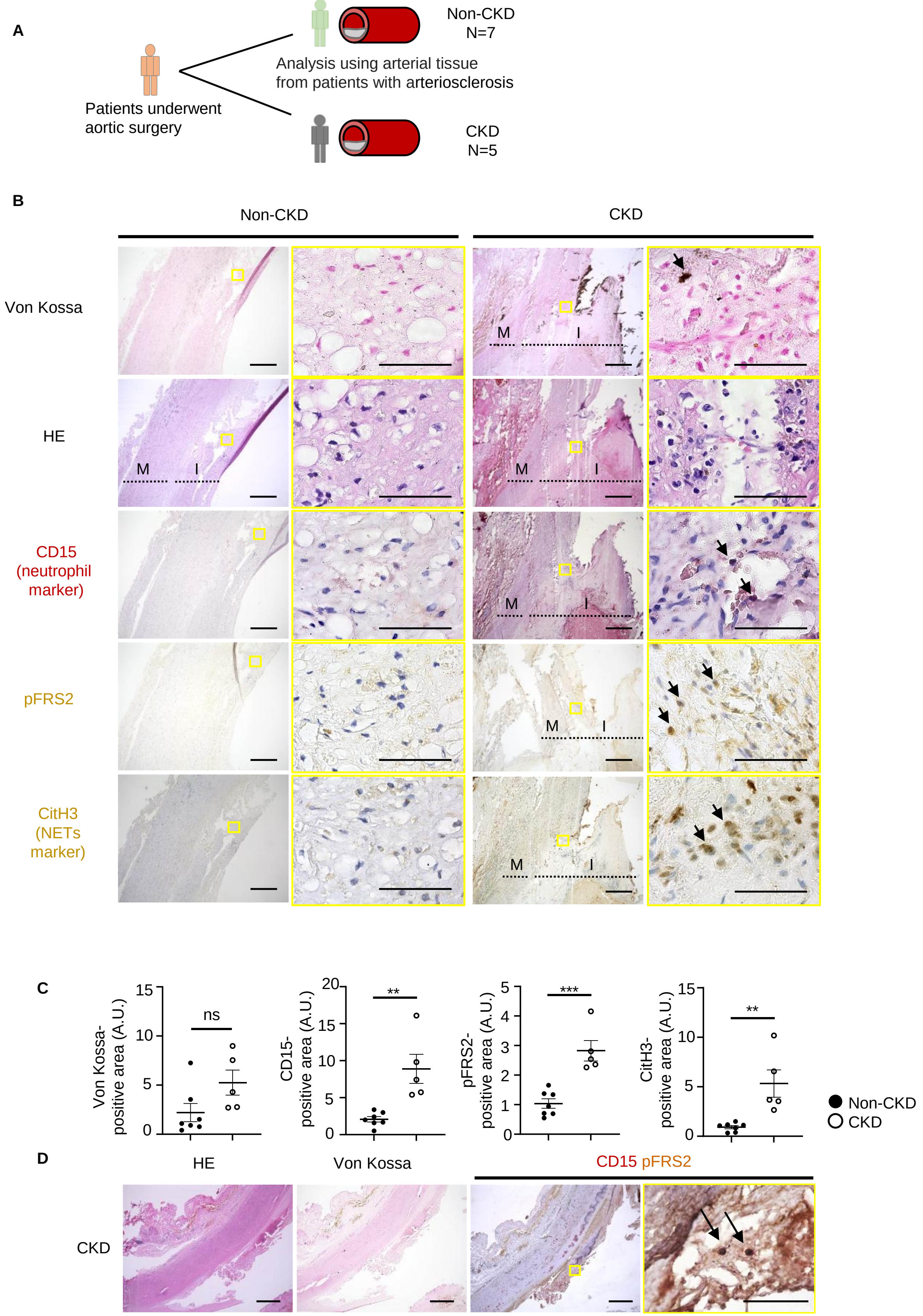

Supplemental Figure 8

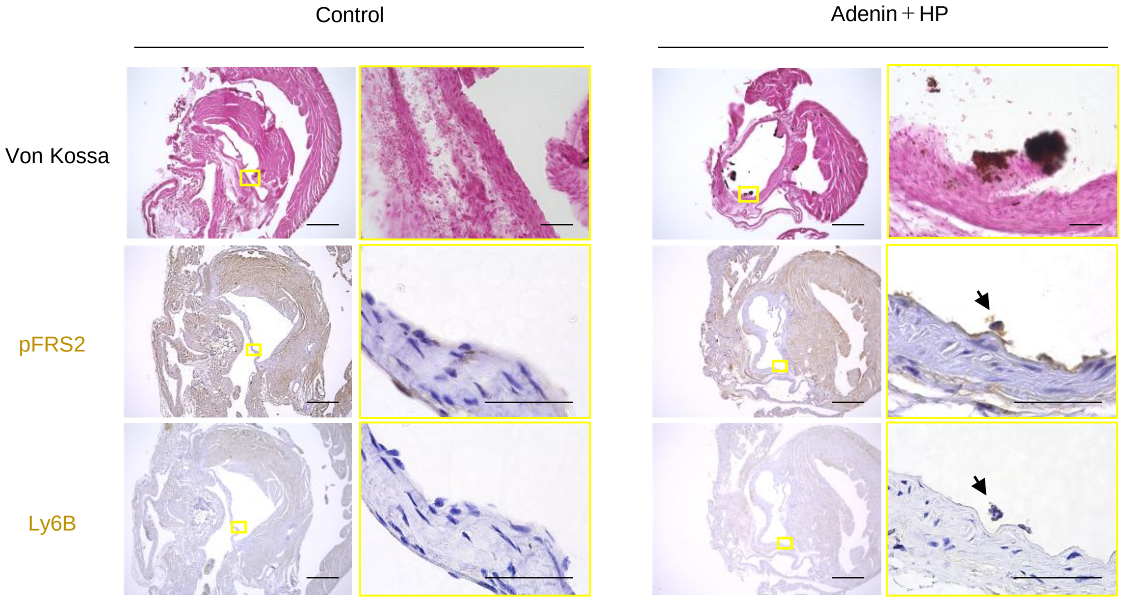

Supplemental Figure 9

A

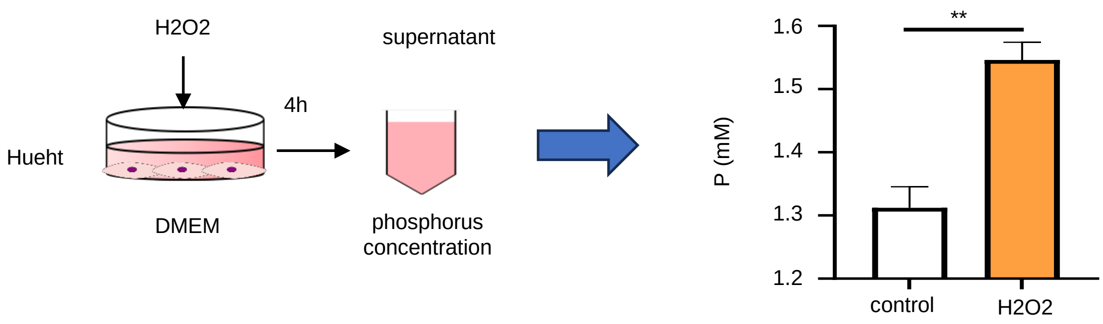
