## Supplemental information for "FGFR1-phosphate sensing and crystal-induced gasdermin D signaling in neutrophils drive vascular calcification in CKD"

**Supplemental Figure Legends**

**Supplemental Figure 1. The response to phosphate among healthy individuals and pre-dialysis CKD patients.**

**(A)** Neutrophils isolated from both HC and pre-dialysis CKD patients were stimulated with various P concentrations. **(B)** The representative images of SYTOX green staining. Scale bars, 200 μm. **(C)** The comparison of quantification of SYTOX-positive area between HC (●) and CKD (❍). n=3 for each group. Data represent mean ± SEM. Statistical analysis was performed using Student’s unpaired t test. *P <0.05, **P <0.01.

**Supplemental Figure 2. Response to elevated phosphate levels in lymphocytes and monocytes.** (**A** and **B**) Lymphocytes (**A**) and monocytes (**B**) were incubated for 6 hours in 2.5% FBS DMEM medium with various concentrations of phosphate buffer, and stained with SYTOX green (green). Right graphs show the quantification of SYTOX-positive area. n=4 for each group. Statistical analysis was performed using one-way ANOVA with post hoc Dunnett’s multiple-comparison test. ns: not signiﬁcant. Data represent mean ± SEM. Scale bars, 200 μm.

**Supplemental Figure 3. The cytotoxicity of FGFR1 inhibitor in human neutrophils using the cell counting kit8.** The FGFR1 inhibitor was administered at concentrations ranging from 0 to 80 μM, with H₂O₂ serving as the positive control.

**Supplemental Figure 4. The viability of HL-60 cells through transfection.** The graph indicates the viability of si control HL-60 cells (green) and si FGFR1 HL-60 cells (orange) through transfection using the trypan blue.

**Supplemental Figure 5. Neutrophils treated with varying concentrations of phosphate and calcium without FBS.** (**A** and **B**) Neutrophils isolated from both HC and CKD patients were stimulated with varying concentrations of phosphate and calcium without FBS. The results show representative images and a comparison of the quantification of SYTOX-positive area between HC (●) and CKD patients (❍). n=5 for each group. (**C** and **D**) Neutrophils derived from HC were incubated for 3 hours in DMEM medium with 2.5% FBS or 2.5% bovine serum albumin (BSA) supplemented with low or high phosphate. The results show representative images and a comparison of the quantification of SYTOX-positive area between 2.5% FBS (■) and 2.5% BSA (□). n=5 for each group. (**E** and **F**) Healthy neutrophils pretreated with 20 μM FGFR1-I (SU5402) were cultured under high phosphate and calcium conditions for 3 hours without FBS. Data represent mean ± SEM. Statistical analysis was performed using one-way ANOVA followed by post hoc Dunnett’s multiple-comparison test (F) or Student’s unpaired t test (B and D). *P <0.05, **P < 0.01, ***P < 0.001, ****P < 0.0001. ns: not signiﬁcant. The insets are higher magniﬁcation images of the black and yellow boxes. Scale bars: 200 μm (A, C, and E).

**Supplemental Figure 6 NET-induced endothelial injury caused by high phosphate and calcium phosphate crystals in vitro.**

(**A**) In the phosphorus-dependent model, neutrophils pretreated with either a 20 μM FGFR1 inhibitor or a 200 μM PAD4 inhibitor were incubated for 6 hours in 2.5% FBS DMEM supplemented with low or high phosphate on a 0.4-μm culture insert. In the calcium phosphate crystals-dependent model, neutrophils pretreated with a 40 μM GSDMD inhibitor were incubated for 3 hours in DMEM without FBS supplemented with low or high phosphorus and calcium on a culture insert. HuEht cells stimulated by NETs composition were stained with calcein-AM (green) and propidium iodide (PI; red). (**B**) The panels show the representative fluorescent images of calcein-AM/PI stained HuEht cells treated with each inhibitor in the phosphorus-dependent model. Right graph is cytotoxicity as a percentage of the calcein-positive area relative to the sum of the calcein- and PI-positive area. n=6 for each group. (**C**) Representative images and cytotoxicity of HuEht cells treated with each inhibitor in the calcium phosphate crystals-dependent model. n=6 for each group. (**D**) mRNA expressions of IL-1β, IFIT1, CXCL1, and IL-8 assessed by PCR (n=3 for each group). Total RNA was extracted from HuEht cells in the calcium phosphate crystals-dependent model. Data represent mean ± SEM. Statistical analysis was performed using one-way ANOVA followed by post hoc Tukey’s test (B, C and D). *P <0.05, **P < 0.01, ***P < 0.001, ****P <0.0001. Scale bars: 200 μm (B and C).

**Supplemental Figure 7. The expression of pFRS2 in neutrophils within intimal calcification.** (**A**) Illustration of the experimental design. Patients with arteriosclerosis who underwent aortic surgery were divided into CKD and non-CKD groups. Arterial tissue from these patients was then histologically analyzed for arteriosclerosis: CKD group (N=5) and non-CKD group (N=7). (**B**) The panels show the representative images of von Kossa, Hematoxylin-eosin (HE), CD15 (red) -, pFRS2 (brown) -, and CitH3 (brown) -stained sections of human aorta. The black arrows indicate calcium phosphate crystal deposit, CD15-, pFRS2-, and CitH3-positive cells, respectively. Scale bars: 500 μm (low magnification) and 50 μm (high magnification). M: media. I: intima. (**C**) The quantification of von Kossa-, CD15-, pFRS2-, and CitH3-positive area with statistical analysis performed using Student’s unpaired t test between non-CKD (●) and CKD patients (❍). **P < 0.01, ***P < 0.001. ns: not signiﬁcant. (**D**) Representative images of CD15(red) and pFRS2 (brown) immunostaining of the aorta in CKD group. The black arrow indicates CD15 and pFRS2 double-positive cells. Scale bars: 250 μm (low magnification) and 50 μm (high magnification). The insets are higher magniﬁcation images of the yellow boxes.

**Supplemental Figure 8. The expression of pFRS2 in neutrophils within intimal calcification in mice.** The representative images of Von Kossa, pFRS2, and Ly6B-stained sections of the aortic root in CKD calcification model induced by adenine and high phosphorus diet food. The black arrows (→) indicate pFRS2- and Ly6B- positive cells, respectively. Scale bars: 500 μm (low magnification) and 50 μm (high magnification).

**Supplemental Figure 9. Phosphorus concentration changes induced by cell death.** (**A**) HuEht cells treated with 5 mM hydrogen peroxide (H₂O₂) were incubated in DMEM medium for 4 hours to induce necrosis. Subsequently, the concentrations of phosphate in the supernatant were measured. The graphs show concentrations of phosphate, with statistical analysis performed using Student’s t test. n=4 for each group. Data represent mean ± SEM. **P <0.01.

**Supplemental Tables**

**Supplemental Table 1. Characteristics of CKD patients for** **the experiment on the addition of high levels of phosphorus to neutrophils.**

|  | Pre-dialysis  CKD patient  (N=4) | HC  (N=4) |
| --- | --- | --- |
| Age | 55 (44-64.5) | 44 (42-44) |
| Sex (M/F) | (2/2) | (2/2) |
| Cause of CKD, n |  |  |
| Diabetic nephropathy | 2 |  |
| Nephrosclerosis | 1 |  |
| Glomerulonephritis | 1 |  |
| Hemoglobin (g/dL) | 11.1 (10.7-11.5) |  |
| Albumin (g/dL) | 2.3 (1.9-3.0) |  |
| BUN (mg/dL) | 52 (33-77) |  |
| Creatinine (mg/dL) | 4.2 (3.0-6.1) |  |
| eGFR (ml/min/1.73m^2^) | 12.3 (6.7-25.0) |  |
| Calcium (mg/dL) | 9.2 (8.4-10.0) |  |
| Phosphorus (mg/dL) | 4.2 (3.6-5.2) |  |
| HbA1c (%) | 6.3(5.6-7.1) |  |

Values for continuous variables are presented as medians (interquartile range).

BUN indicates blood urea nitrogen; CKD, chronic kidney disease; F, female and M, male.

**Supplemental Table 2. Characteristics of arteriosclerosis patients for aorta histology.**

|  | Non- CKD (N=7) | CKD patient (N=5) |
| --- | --- | --- |
| Age | 76 (65-80) | 70 (69-76) |
| Sex (M/F) | (2/5) | (4/1) |
| Cause of CKD (n) |  |  |
| Nephrosclerosis |  | 2 |
| Polycysic kidney disease |  | 1 |
| Glomerulonephritis |  | 2 |
| Hemoglobin (g/dL) | 12.7 (12.1-13.0) | 11.6 (11.5-12.9) |
| Albumin (g/dL) | 3.7 (3.5-4.1) | 4.1 (4.0-4.2) |
| BUN (mg/dL) | 18 (16-21) | 47 (32-48) |
| Creatinine (mg/dL) | 0.7 (0.6-0.9) | 2.0 (1.8-3.8) |
| eGFR (ml/min/1.73m^2^) | 66.9 (60.1-72.4) | 7.3 (10.1-28.3) |
| Calcium (mg/dL) | 9.0 (8.8-9.2) | 9.1 (9.0-9.2) |
| Phosphorus (mg/dL) | 3.3 (3.1-3.7) | 4.2 (3.4-4.9) |
| HbA1c (%) | 5.6 (5.5-5.8) | 5.6 (5.2-6.0) |

Values for continuous variables are presented as medians (interquartile range).

Abbreviations are explained in Supplemental Table 1.
